# The Relationship Between Genome Size and Metabolic Rate in Extant Vertebrates

**DOI:** 10.1101/659094

**Authors:** Jacob D. Gardner, Michel Laurin, Chris L. Organ

**Author notes:** Corresponding author’s e-mail addresses (J. D. Gardner), (M. Laurin), (C. L. Organ).

## Abstract

Genome size has long been hypothesized to affect metabolic rate in various groups of animals. The mechanism behind this proposed association is the nucleotypic effect, in which large nucleus and cell sizes influence cellular metabolism through surface area-to-volume ratios. Here, we provide a review of the recent literature on the relationship between genome size and metabolic rate. We also conduct an analysis using phylogenetic comparative methods and a large sample of extant vertebrates. We find no evidence that the effect of genome size improves upon models in explaining metabolic rate variation. Not surprisingly, our results show a strong positive relationship between metabolic rate and body mass, as well as a substantial difference in metabolic rate between endothermic and ectothermic vertebrates, controlling for body mass. The presence of endothermy can also explain elevated rate shifts in metabolic rate whereas genome size cannot. We further find no evidence for a punctuated model of evolution for metabolic rate. Our results do not rule out the possibility that genome size affects cellular physiology in some tissues, but they are consistent with previous research suggesting little support for a direct functional connection between genome size and basal metabolic rate in extant vertebrates.

## Introduction

Genome size (haploid C-value) can, theoretically, explain various physiological and developmental traits, including metabolic rate [1,2]. Gregory [2] gave several theoretical reasons why genome size should correlate with nucleus size, cell size, cell division rate, and hence, organismal developmental rate. Many studies have tested the relationship between genome size and metabolic rate using a variety of different methods and taxonomic samples (many reviewed by Gregory [3]). The current literature points to a possible difference in this relationship between endothermic and ectothermic species. The evidence generally suggests that a relationship might exist in birds and mammals but not in ectothermic groups, specifically actinopterygians and lissamphibians [3]. The evidence, however, on the relationship between genome size and metabolic rate in birds is conflicting among studies [4–8]; this is likely due to differences in sample size and methodology (e.g., ordinary least-squares regression, regression of extracted residuals from body mass correlations, and the use of phylogenetic comparative methods). Previous research also suggests that differences in evolutionary mode (gradual vs punctuated change) between genome size and metabolic rate may also diminish the power to detect a correlation between the two [7,9].

The objectives of this article are to 1) review the current literature on the relationship between genome size and metabolic rate, and 2) use Bayesian phylogenetic comparative methods to analyze these traits for a large sample of vertebrate species. Specifically, we will test for a correlation between basal metabolic rate and genome size, accounting for body mass and variable rates of evolution. We will also test for a difference in this relationship between endothermic and ectothermic species, as has been previously suggested [3]. Lastly, we will evaluate the evolutionary mode of basal metabolic rate by testing if the rate of trait evolution is correlated with the number of speciation events.

We begin by reviewing previous studies on the relationship between metabolic rate and genome size (Table 1) in amniotes, lissamphibians, and other vertebrates (mainly actinopterygians). Herein, we abbreviate basal, resting, and standard metabolic rate into BMR, RMR, and SMR, respectively. We use “metabolic rate” when discussing the trait generally.

**Table 1.** Summary of previous research testing for an effect of genome size on metabolic rate.

| Authors (Year) | Taxa | Method | Result | Reference |
| --- | --- | --- | --- | --- |
| Vinogradov (1995) | Birds and mammals | Multiple regression of extracted residuals | Negative effect in mammals but not in birds. | [4] |
| Smith et al. (2017) | Bats | Simple regression of extracted residuals. Independent contrasts not used for BMR correlations. | Evidence for negative effect only at species level w/o mass correction. No evidence otherwise. | [10] |
| Vinogradov (1997) | Birds | Phylogenetic independent contrasts (PIC) of extracted residuals | Negative effect | [5] |
| Gregory (2002) | Birds | Simple regression of extracted residuals | Negative effect | [6] |
| Waltari and Edwards (2002) | Vertebrates | PIC | No effect globally but negative for amniotes and archosaurs. | [7] |
| Kozłowski et al. (2003) | Birds and mammals | Simple regression of genome size and BMR slope coefficients | Negative effect | [14] |
| Uyeda et al. (2017) | Vertebrates | Multiple regression using a phylogenetic Ornstein-Uhlenbeck model | Genome size effect not included in final model | [9] |
| Ji and DeWoody (2017) | Birds | PIC | No effect | [8] |
| Licht and Lowcock (1991) | Urodeles | Simple regression at different temperatures | Slight negative effect only at 25°C | [34] |
| Gregory (2003) | Lissamphibians | Simple regression of extracted residuals | Negative overall but not within anurans and urodeles | [35] |
| Hardie and Herbert (2004) | Actinopterygians | Simple regression | No effect at the family level | [40] |
| Smith and Gregory (2009) | Actinopterygians | Simple regression of extracted residuals | No effect | [41] |

## Amniotes

### Correlating Genome Size and Metabolic Rate

Vinogradov [4] studied the relationship between RMR, body mass, and genome size in birds and mammals. He found strong evidence for a relationship between RMR and body mass with the latter explaining 70 to 92% of the variance in RMR, depending on the taxon (birds or mammals) and the taxonomic level (data were grouped within species, genera, families, and orders). No evidence for a correlation was found when he analyzed the raw data. However, when using the residuals of a regression model between RMR and body mass, he found evidence for a negative effect of genome size on mass-corrected RMR. The correlation was significant only in mammals, in which genome size explained 7 (at the species level) to 20% (at the order level) of the variance in RMR. Vinogradov [4] attributed the lack of evidence for a relationship in birds to the stronger relationship between RMR and body mass, which, consequently, leaves less room for the effect of genome size. This, combined with the low variability in genome size in birds, means that greater measurement precision is required to detect an effect; indeed, he casts doubt on the precision of some genome size measurements from the literature. Low genome size variability was also discussed by Smith et al. [10] when they found no evidence for a correlation between genome size and metabolic rate in bats after accounting for body mass. The study by Vinogradov [4], however, incorporated only rough phylogenetic information where the data were treated as statistically independent at each taxonomic level. Common statistical tests assume that the residuals associated with the data are independently structured. However, residual independence of interspecies data cannot be assumed due to common ancestry, even when analyzed at different taxonomic levels [11]. Also, the higher-level taxonomic groups (e.g., families and orders) that Vinogradov [4] used may not be monophyletic, which raises problems in comparative biology generally [12,13]. We extend this concern for all subsequently mentioned literature that found evidence for a correlation and possessed similar methodologies.

Vinogradov [5] reanalyzed the data on passeriform birds using phylogenetic independent contrasts, which creates independently structured contrasts from the trait data. In all cases, instead of regressing raw values, he regressed both RMR and genome size against body mass first (all variables log-transformed) to extract residuals. Mass-corrected RMR and genome size were then regressed against one another. Even though only 11 nominal species were included (representing seven genera and four families), he found statistically significant associations in all analyses. Gregory [6] used a methodology similar to Vinogradov [5]—regressed extracted residuals from a correlation with body mass—on an expanded dataset of birds (50 species) and found evidence for a relationship between genome size and RMR. The larger sample size in this study suggests that a relationship may exist between genome size and metabolic rate in birds when a sufficient statistical power is achieved. However, this study did not account for the residual non-independence of interspecies trait data.

Waltari and Edwards [7] analyzed a dataset comprised of 91 vertebrate taxa, emphasizing archosaurs (one crocodilian and 19 bird taxa), using phylogenetic independent contrasts. They found little to no evidence for a relationship among vertebrates globally, but they found an inverse relationship between BMR and genome size in amniotes and in archosaurs. Waltari and Edwards [7] also suggest a punctuated evolutionary model for genome size while a more gradual one for BMR—although they do not test for it directly. The authors suggest that differences in evolutionary mode may reduce any potential association between the two traits. According to Waltari and Edwards [7], it is too simplistic to expect a simple, tight relationship between genome size and metabolic rate. Waltari and Edwards [7] also tested for a correlation between intron length and BMR—higher metabolic rates may lead to genome contraction by shortening intron lengths— but they found no evidence for a relationship.

Kozlowski et al. [14] developed a new model based on theoretical considerations to study the relationship among BMR, body mass, and cell size, which is itself correlated with genome size. Their model suggests that genome size should have a strong effect on BMR. They test this model by correlating the allometric slope coefficients of genome size and BMR from regressions with body mass and found evidence for an inverse relationship. However, their analyses did not use phylogenetic comparative methods.

Uyeda et al. [9] studied the effect of genome size on SMR and BMR (among other questions) in a large number of vertebrate taxa that emphasized amniotes (n = 857, though only 318 had data on genome size; the others were estimated using a Brownian motion process on a time tree using an MCMC procedure). They fit several Ornstein-Uhlenbeck phylogenetic models (with and without genome size) to explain the evolution of SMR and BMR. The best model according to AIC model selection did not include genome size—although a model including genome size found some evidence for a small negative effect. The authors concluded that genome size may well have an effect, but that metabolic rate may be affected by other variables and that the complex evolutionary pattern of genome size may obscure this relationship. Uyeda et al. did not test for a difference in the effect of genome size between endothermic and ectothermic species, as has been previously suggested [3].

### The Effect of Powered Flight on Genome Size

Ji and DeWoody [8] found no evidence for a relationship between BMR and genome size using phylogenetic independent contrasts on a sample of 13 bird taxa. They hypothesize, however, that BMR may contribute to shaping the transposable element landscape in birds and play a role in driving the rate of insertions and deletions (indels). They argue that higher BMRs are associated with faster cell cycles and, therefore, result in more replication-dependent mutations. Kapusta et al. [15] corroborate this hypothesis by finding that volant birds and mammals (i.e., bats) exhibited a higher rate of indels despite having relatively low variation in genome size. Their ‘accordion model’ posits that transposable element accumulation is quickly counteracted by large segmental deletions; volant birds and mammals have smaller genomes but more dynamic gains and losses, specifically more extensive loss. Flightless birds were found to have considerably lower midsize deletion rates and less-recycled transposable elements (i.e., fewer DNA losses and gains through the accordion process). This supports the hypothesis that genome contraction and metabolic rate are connected; however, this does not necessarily suggest that small genomes were an adaptation for powered flight. Kapusta et al. [15] also found that larger species have less-dynamic genomes compared to smaller species within the same order; body size is strongly associated with metabolic rate.

Many other studies support the hypothesis that higher metabolisms are associated with genome contraction. A study comparing genome sizes among birds found that volant birds have smaller genomes on average than non-volant birds [16]. Zhang and Edwards [17] also demonstrate that birds and bats have shorter introns on average than non-volant vertebrates. A study on megabats found that they exhibit less genome size variation and, often times, smaller genomes than microbats despite having larger body sizes [18]. Indeed, vertebrate-wide studies have not found evidence for a relationship between genome size and body size [3,19]. This is inconsistent with models proposing effective population size as the dominant factor in driving genome size evolution [20]; small populations are governed by non-adaptive processes (e.g., genetic drift), which can lead to the retention of duplicate genes and transposable elements. Instead of correlating genome size with metabolic rate directly, numerous studies correlate the former with multiple flight efficiency variables, such as flight muscle size, heart index, wing area, and wing loading index. These studies found good evidence for a relationship between these variables and genome size in birds [21,22] and bats [10] but not within hummingbirds [23]. These studies lend further support for a link between high metabolic activity and genome size, but whether small genomes were an adaptation for the metabolic demands of powered flight remains uncertain.

The fossil record has also contributed many insights into the hypothesis that the metabolic demands of powered flight selected for smaller genomes [24,25]. Organ et al. [24] leveraged the statistical association between genome size and osteocyte lacunae size to predict the genome sizes for multiple dinosaur species. They found that birds inherited their small genomes from earlier flightless theropods—a group of mostly carnivorous dinosaurs that also includes taxa like *Tyrannosaurus rex* and *Velociraptor*. Organ et al. [25] used the same approach to predict the genome sizes of pterosaurs—an extinct group of Mesozoic reptiles (closely related to dinosaurs) that were the first vertebrates to evolve powered flight. They found that pterosaurs also had relatively small genomes, providing further support for the hypothesis that powered flight is associated with genome contraction. Organ et al. [25] also found support for a proportional model of genome size evolution, where larger genomes evolve faster than smaller ones [26]. It was, therefore, unlikely for the common ancestor of dinosaurs and pterosaurs to have had a small genome because the larger genomes of ornithischian dinosaurs (e.g., *Triceratops* and *Stegosaurus*) were unlikely to have evolved from a small genome. It is more likely that small genomes evolved independently in pterosaurs and theropod dinosaurs. Altogether, this research suggests that small genomes were necessary for the evolution of powered flight and that the metabolic demands of it sustained genome size contraction.

### Genome Size and Osteocyte Lacunae Size

A relationship may exist between osteocyte lacunae size and metabolic rate given that genome size is correlated with the former [24,27]. However, a recent study on archosaurs did not find a significant relationship between osteocyte lacunae size and RMR [25; Legendre, pers. Comm. January 27, 2019]. This suggests that there is no relationship between genome size and RMR in Legendre et al.’s [28] dataset. In fact, the link between cell size and RMR is more direct and stronger than between genome size and RMR, as suggested both by theoretical considerations [2] and empirical findings. For instance, Starostova et al. [29] found a significant relationship between SMR and cell size, but not with genome size, in a sample of 14 eyelid gecko taxa (analyzed with and without phylogenetic comparative methods). The lack of a relationship between osteocyte lacunae size and RMR found by Legendre et al. [28] might reflect the relatively small dataset (14 extant terminal taxa, including one amphibian and 13 amniotes, with RMR data; 14 extinct terminal taxa were also included, but only to infer their RMR). Note that this study used a different comparative method, phylogenetic eigenvector maps [30]. Another possible explanation for this negative result is provided by Grunmeier and D’Emic [31], who found no correlation between BMR and the volume of bird femoral osteocyte lacunae in bone derived from static osteogenesis, whereas such a relationship was found with lacunae located in bone developed through dynamic osteogenesis. A similar study, which excluded osteocyte lacunae formed from static bone formation, also found a weak correlation between osteocyte lacunae volume and mass-specific BMR [32]. Legendre et al. [28] did not specify if the lacunae that they measured were located in bone issued from dynamic or static osteogenesis, so no firm conclusions can be drawn on this point. However, these results suggest that other bone histological data beyond osteocyte lacunae size (e.g., vascularization, tissue type, etc.) is needed to evaluate the relationship between bone growth and metabolism.

Beyond osteocytes, metabolic rate was found to scale differently depending on the cell type [33]. Mass-specific BMR in birds was found to decrease with an increase in the size of skin and kidney cells, chondrocytes (cartilage), enterocytes (small intestine), and erythrocytes (red blood cells), but was found to increase with liver cell size. An inverse correlation between RMR and red blood cell size was previously found in birds [6]. The negative association between BMR and the size of, at least, five different cell types is consistent with a prediction from the optimal cell size hypothesis that large-bodied organisms reduce energetic waste by increasing cell size and maintaining operational cell membranes [33].

### Summary

Studies generally support a link between genome size and metabolic rate in amniotes [4–7,14], except in a few cases [4,8,10]. Differences in support among these studies may be due to methodology, sample size, and taxonomic focus (denser sampling of one clade vs broader sample). The study with the largest sample size did not find support for including genome size in their final model [9]. However, they did not test for a difference in the effect of genome size between endothermic and ectothermic species; this could diminish a possible effect in birds and mammals (as previously found) if there is no evidence for an effect in ectothermic taxa.

## Lissamphibians

The evidence for a relationship between genome size and metabolism is hardly more convincing for lissamphibians. Licht and Lowcock [34] tested for a relationship between genome size and SMR on urodeles at four temperatures (5, 15, 20, and 25° C) without phylogenetic comparative methods. In a series of analyses using all the data available (21 to 39 species because not all data were available for all taxa at all temperatures), they found a significant relationship at 15 and 25°C. However, after removing two outliers (*Necturus maculosus* and *Amphiuma means*) the relationship remained significant only at 25°C. Given that 25°C is close to the lethal limit for many urodele taxa, Licht and Lowcock [34] concluded that “the prediction that nuclear and cell size, mediated via genome size, will have the presumed effects for metabolism in salamanders under a normal thermal range of activity does not appear substantiated.” The largest study on lissamphibians corroborated these results by finding a negative effect of genome size on RMR overall but not within anurans and urodeles [35].

A study by Hermaniuk et al. [36] on diploid and triploid edible frogs (*Pelophylax esculentus*) illustrates the complexity of this issue. *P. esculentus* is a natural hybrid of *Pelophylax lessonae* (genotype LL) and *Pelophylax ridibundus* (RR). Hermaniuk et al. [36] measured the SMR of tadpoles and froglets of LLR triploids and LR diploids to test the hypothesis that the larger genome, nucleus, and cell size of the triploids resulted in a lower SMR. After checking for possible gene dosage effects and the size of the metabolically most active organs, they concluded that triploid tadpoles indeed had a lower SMR than diploids (p = 0.036), but that no such effect was detected with the froglets (p = 0.255). They explained these contrasting results by the lower concentration and diffusion rate of O_2_ in water compared to air (33 times less and 3 x 10^−5^ times slower), which means that O_2_ concentration might be more limiting for large cells in aquatic organisms compared with terrestrial species. This explanation is testable, for instance, using a large sample of aquatic and terrestrial lissamphibian taxa, but such a study has not yet been conducted, as far as we know.

The latest study on lissamphibians found a direct relationship between developmental time (from zygote to birth in taxa with direct development; from zygote to metamorphosis in other taxa) and genome size in anurans, but not in urodeles [37]. However, the direction of this relationship could not be determined; it is unclear if developmental time constrains genome size or the reverse. No link was found between genome size and developmental complexity, contrary to previous suggestions [38]. A possible link with basal metabolism was not investigated, perhaps because BMR data were not available for a sufficient number of lissamphibian taxa, because BMR does not vary enough within this group, or because previous studies raised doubts about the presence of such a relationship [35].

Previous studies and reviews on the relationship between genome size and metabolism question the role of large genomes driving lower metabolic rates in lissamphibians, particularly in aestivating species [3,39]. Gregory [39] demonstrates that a large genome is not necessary for maintaining an aestivating lifestyle. For example, two aestivating frog species, *Scaphiopus couchii* and *Pyxicephalus adspersus*, can lower their metabolic rates to the level of the urodele *Siren intermedia* despite having smaller genomes [3,39].

The above studies, overall, suggest that there is little evidence for a relationship between genome size and metabolic rate within anurans and urodeles.

## Other vertebrates

The relationship between genome size and metabolic rate is poorly studied in other vertebrates with a few notable exceptions. Hardie and Hebert [40] found no relationship between genome size and standard or routine metabolic rate (i.e., average rate when undergoing a defined type of activity) in actinopterygians. Their sample size was modest (24 and 37 nominal families for standard and routine metabolic rates) but several interesting results illustrate the numerous constraints that may influence genome size evolution. For instance, they found that marine and catadromous actinopterygians had a significantly smaller genome (1.77 pg) than freshwater and anadromous relatives (2.81 pg; p < 0.0001). Genome size was estimated for the whole nucleus, rather than for a haploid genome, which raises the possibility that polyploidy may have influenced the results. The authors performed a second test on diploid taxa and the result remained highly significant (p < 0.0001) with larger genomes of freshwater and anadromous taxa (2.32 pg) compared with marine and catadromous relatives. Hardie and Hebert [40] also found a strong positive relationship between genome size and egg size in actinopterygians (r^2^ = 0.41, p < 0.005 for a linear regression model of 18 nominal orders based on 88 species). Smith and Gregory [41] also found no evidence for a relationship between genome size and routine metabolic rate using simple linear regressions on the residuals extracted from body mass correlations. No evidence was found at any taxonomic level and regardless of whether or not chondrosteans and polyploids were included. Maciak et al. [42] assessed the effect of ploidy and other factors on metabolic rate in the *Cobitis taenia* hybrid complex, which includes diploid and triploid hybrids. They found that ploidy level explains 17% of the variation within SMR (p = 0.03). This does not contradict previous studies given their broader taxonomic samples compared to Maciak et al.’s focus on one genus [40,41]. Furthermore, Maciak et al. [42] cite previous studies that failed to find differences in SMR in di- and triploids of other teleost hybrids. These results suggest that genome size influences various biological attributes in actinopterygians but not necessarily metabolic rate.

## Summary of previous studies

Altogether, previous research suggests that a negative effect of genome size on metabolic rate may exist in amniotes but not in actinopterygians and lissamphibians (Table 1). Studies on actinopterygians, lissamphibians, and mammals (except within bats) are fairly consistent in recovering a negative effect, but studies on birds are less consistent. This is likely due to differences in methodology, sample size, and taxonomic focus. Some studies use phylogenetic comparative methods while others analyze their data at different taxonomic levels. Many studies correlate the residuals extracted from body mass regression models while others use a multiple regression framework. It is also possible that limited genome size variation (e.g., birds and bats), the high explanatory power of body size, unaccounted for life-history variables, and/or differences in evolutionary mode can explain differences in the estimated effect of genome size among studies [4,7,9,10]. The largest study so far, incorporating vertebrate-wide data, did not include genome size in its final model, suggesting that genome size is not a sufficiently influential variable in explaining metabolic rate variation after accounting for body mass [9]. However, no study has yet used a large sample size of vertebrates to directly test for a difference in the effect of genome size between endotherms and ectotherms, as has been previously suggested [3]. Moreover, no study has yet accounted for variable rates of evolution under a unified statistical framework.

Here, we test for the effect of genome size on BMR, accounting for body mass, using a large sample of vertebrate species and a recently-developed regression model that allows for variable rates of evolution. We further test for a difference in the effect of genome size on BMR between endothermic and ectothermic species. Lastly, we evaluate the evolutionary mode (gradual vs punctuated) of BMR while accounting for body mass and rate variation.

## Methods

To clarify the relationship between genome size and metabolic rate, we test for an association between the two (accounting for body mass) for a dataset of vertebrates using phylogenetic comparative methods. We used the average BMR (corrected to 20°C), body mass, and haploid genome size (C-value) data from Uyeda et al. [9,43], consisting of 30 actinopterygian, 91 lissamphibian, 34 lepidosaur, 29 bird, and 133 mammal species (317 species total). We used BayesTraits V3 (http://www.evolution.rdg.ac.uk/BayesTraitsV3.0.1/BayesTraitsV3.0.1.html) to create phylogenetic independent contrast (PIC) models and test for a relationship between average BMR and genome size while accounting for body mass. We incorporated a binary “dummy-variable” and its interaction with genome size to test for a difference in the effect of genome size on BMR between endothermic and ectothermic species (0 = ectothermic, 1 = endothermic). To the best of our knowledge, no taxa in our ectotherm sample are homeothermic; given the small overlap in BMR between the two groups, the number of homeothermic ectotherms (if present) is small and unlikely to substantially affect our results. We then used a recently-developed variable rates regression model to test for variable rates in BMR evolution while accounting for the effects of body mass, genome size, and presence of endothermy [44,45]. The variable rates regression model detects evolutionary rate shifts in the unexplained residual variance of a given regression model; branch- and clade-specific rate shifts are proposed using a Bayesian reversible jump Markov-chain Monte Carlo (RJMCMC) procedure, which reduces the number of parameters to those only supported by the data. Baker et al. [44,45] argue for ruling rate shifts as evidence for positive selection when a positive rate shift (r > 1) is observed in at least 95% of the posterior distribution of variable rates models. They also argue that positive selection in a trait can be explained by another trait if adding an additional covariate reduces the number of “significant” rate shifts (r > 1 in ≥ 95% posterior distribution). Although this approach can detect highly positive rate shifts post hoc, we hypothesize *a priori* that we will observe a rate shift in BMR along the branches leading to birds and mammals. We further hypothesize that these rate shifts will diminish when we include the presence of endothermy in the model. We also test if adding genome size as a covariate explains rate variation in BMR, accounting for body mass.

We used Bayes factors (BF) to compare the variable rates regression models with the original uniform-rate PIC models, where BF > 2 is considered good evidence in favor of the model with the higher log marginal likelihood. We then used a Bayesian Information Criterion (BIC) to compare regression models among one another. BIC compares the log likelihoods among a selection of models while penalizing by the number of parameters. The model with the lowest BIC value is the most supported model with a sufficient number of parameters to explain the data. Models with a difference in BIC of less than two cannot be rejected with statistical confidence.

To determine the mode of evolution in BMR (gradual vs punctuated change), we conducted two tests. For the first, we regressed the path lengths (root to tip lengths) of each species against the net number of nodes (speciation events, conceptualized as cladogenesis, for the purposes of this analysis) along each path length [46,47]. Path lengths were obtained from a tree in which each branch is scaled by the rate of BMR evolution (after applying the variable rates model to BMR). Evidence for an effect of node count on rate-scaled path lengths is consistent with the hypothesis of punctuated evolution—lineages that speciated more frequently exhibited more BMR evolution. For the second test, we included node count as an additional explanatory variable in our final model previously chosen by BIC model selection. This analysis tests if the observed variance in BMR was influenced directionally by the net number of speciation events, either positively or negatively. In other words, we test if BMR tended to increase or decrease along lineages that speciated more frequently. As with all comparative analyses, these two tests for punctuated trait evolution assume an unbiased sample; however, our taxonomic sample is highly disproportionate, resulting in under- and over-estimated node counts when extant diversity is under- and over-represented. By randomly down-sampling our full dataset, we can better approximate the net number of speciation events. To verify our punctuation test results, we excluded the actinopterygians and randomly down-sampled the tetrapod dataset to approximately reflect extant diversity (about 7147 anurans, 738 urodeles, 10418 lepidosaurs, 6399 mammals, and 10966 birds, based on estimates from amphibiaweb.org, reptile-database.org, mammaldiversity.org, and birdlife.org). We randomly sampled 95 taxa from our full dataset to achieve a similarly proportioned sample, including 19 anurans, two urodeles, 28 lepidosaurs, 17 mammals, and 29 birds. We produced three independently down-sampled datasets and repeated the two tests for each. Note that, in our reanalysis of the second test, we assumed the final model selected for the full dataset.

Twice the proportion of the posterior distribution that crosses 0 for regression parameters was used as a measure of statistical significance—referred to as pMCMC. A pMCMC less than 0.05 was used as our indicator for good evidence of a variable’s effect on BMR. We ran all PIC models for 100 million iterations with a 25% burn-in and sampling every 10,000 iterations. We used a Stepping Stone algorithm to estimate the log marginal likelihood of each model, using 100 stones and sampling every 10,000 iterations [48]. We checked regression model assumptions of normality and equal variance in R (Figures S1-S4 for final model). All variables are natural log-transformed (ln), except path length and node count. For models with multiple independent variables, we tested for multicollinearity using variance inflation factors in R (Table 1 in Supplementary Materials). The likelihood convergence of all model MCMC chains were assessed using Tracer 1.7 [49]. Likelihood trace plots are provided for the final uniform-rate and variable rates regression models in the Supplementary Materials (Figures S5 and S6, respectively).

## Results

Our final model includes a body mass effect and a difference in BMR between endotherms and ectotherms (BIC = 814.64). The final model is supported over a model that includes an effect of genome size on BMR (ΔBIC = 35.17) and one that includes an interaction between genome size and the presence of endothermy (ΔBIC = 32.69). The final model is also supported over a model that includes both genome size and a difference in BMR between endotherms and ectotherms (ΔBIC = 13.56) and a model that only includes body mass (ΔBIC = 17.03). We also find compelling evidence for variable rates of BMR evolution, accounting for body mass and a difference in BMR between endotherms and ectotherms (BF = 122.45). For more information regarding our model selection results, please refer to Tables S2-S4. For our final variable rates regression model, there is strong evidence for a positive relationship between ln BMR and ln body mass, conditioning on a difference in ln BMR between endotherms and ectotherms (pMCMC = 0, slope = 0.73). We also find strong evidence for a difference in ln BMR between endotherms and ectotherms (pMCMC = 0; Figure 1). Endotherms were found to have a median BMR (corrected to 20°C) greater than ectotherms by 71.82 mL O_2_ h^-1^ (e^4.27^ = 71.82), accounting for body mass. All final model estimates are detailed in Table S5.

**Figure 1.**
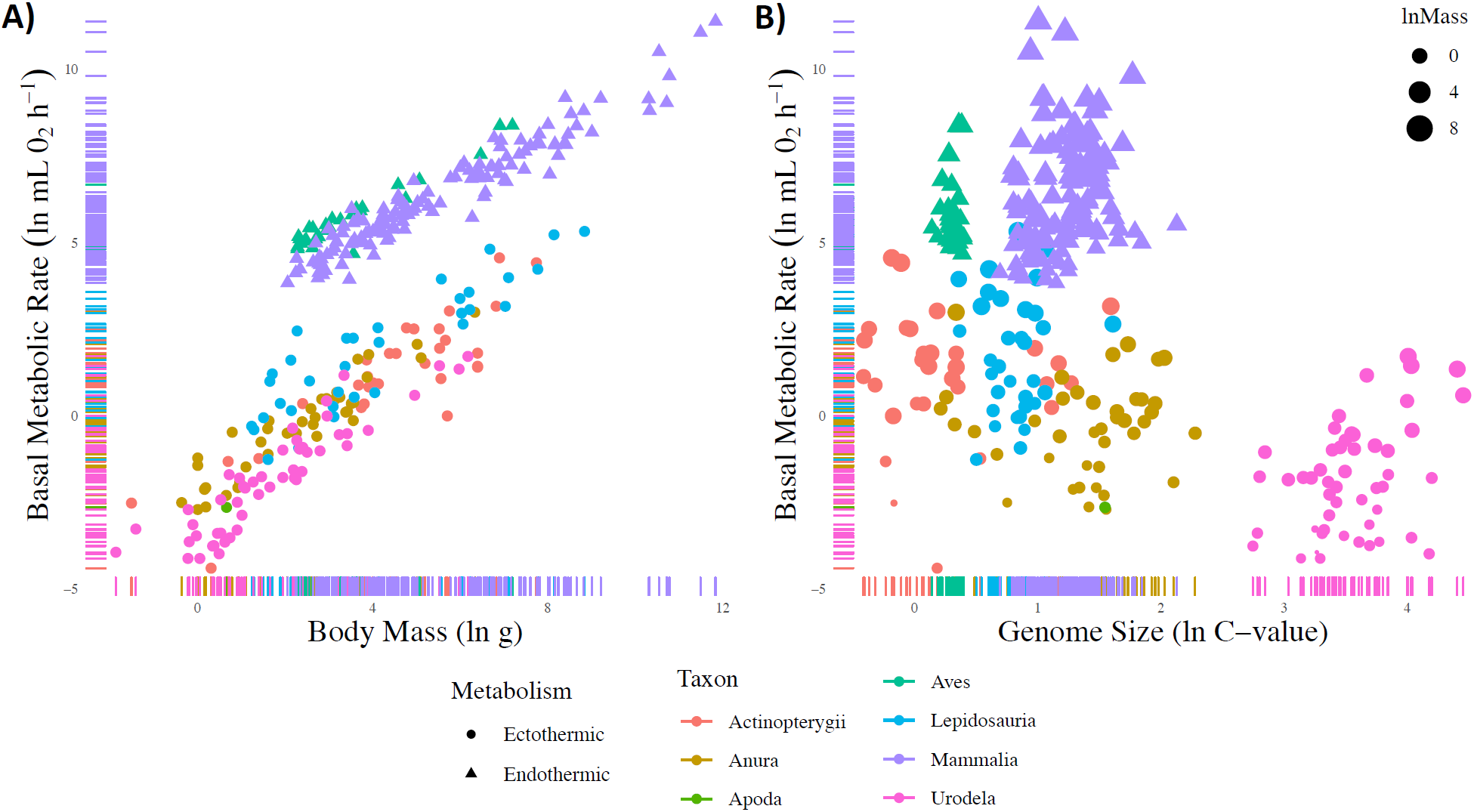
Scatter-plots showing the relationship between the three main studied variables. Circles and triangles represent data from ectothermic and endothermic species, respectively. Data colored by major taxa. A) BMR vs body mass. B) BMR vs genome size. Point size is proportional to body mass.

As hypothesized, the presence of endothermy also explains two positive rate shifts along the branches leading to birds and mammals (Figure 2). A median rate scalar (r) of 15.13 and 13.99 along the branches leading to birds and mammals, respectively, was estimated in a null model that excludes the presence/absence of endothermy. In our final regression model, including the presence/absence of endothermy, the median rate scalars are substantially reduced (r = 1 for both branches). Although these rate shifts are not observed in 95% of the posterior distribution, we argue that they are observed in a sufficiently large proportion of the posterior distribution and result in a substantial reduction in median rate (% posterior for bird and mammal branches = 90.6 and 82.1%, respectively). Moreover, these rate reductions are consistent with our *a priori* hypothesis that the presence of endothermy explains elevated rates along the branches leading to birds and mammals. The number of positive rate shifts in 95% of the posterior distribution of models increased by 14 branches when including the presence/absence of endothermy. However, nearly all of these branches are in close proximity to other rate shifts inferred by the null model and were detected in at least 92% of the null model’s posterior distribution; this suggests that the increase in rate shifts is likely due to model error rather than the presence/absence of endothermy increasing positive rate variation. We also find no evidence that genome size explains rate variation in BMR; adding genome size as a covariate does not reduce the number of positively scaled branches in 95% of the posterior distribution of models (only four different branches are scaled when including genome size, but these can be attributed to model error; i.e., most are sister branches and are observed > 88% of the posterior distribution). This result is consistent with BIC model selection, which did not include genome size in our final model. The majority of highly positive rate shifts (in ≥ 95% posterior distribution) were recovered in the terminal branches of salamanders and frogs as well as a few teleost and squamate species and the large flying fox (*Pteropus vampyrus*). We include a list of all highly positive rate shifts inferred using the final model in the Supplementary Materials (Table S6).

**Figure 2.**
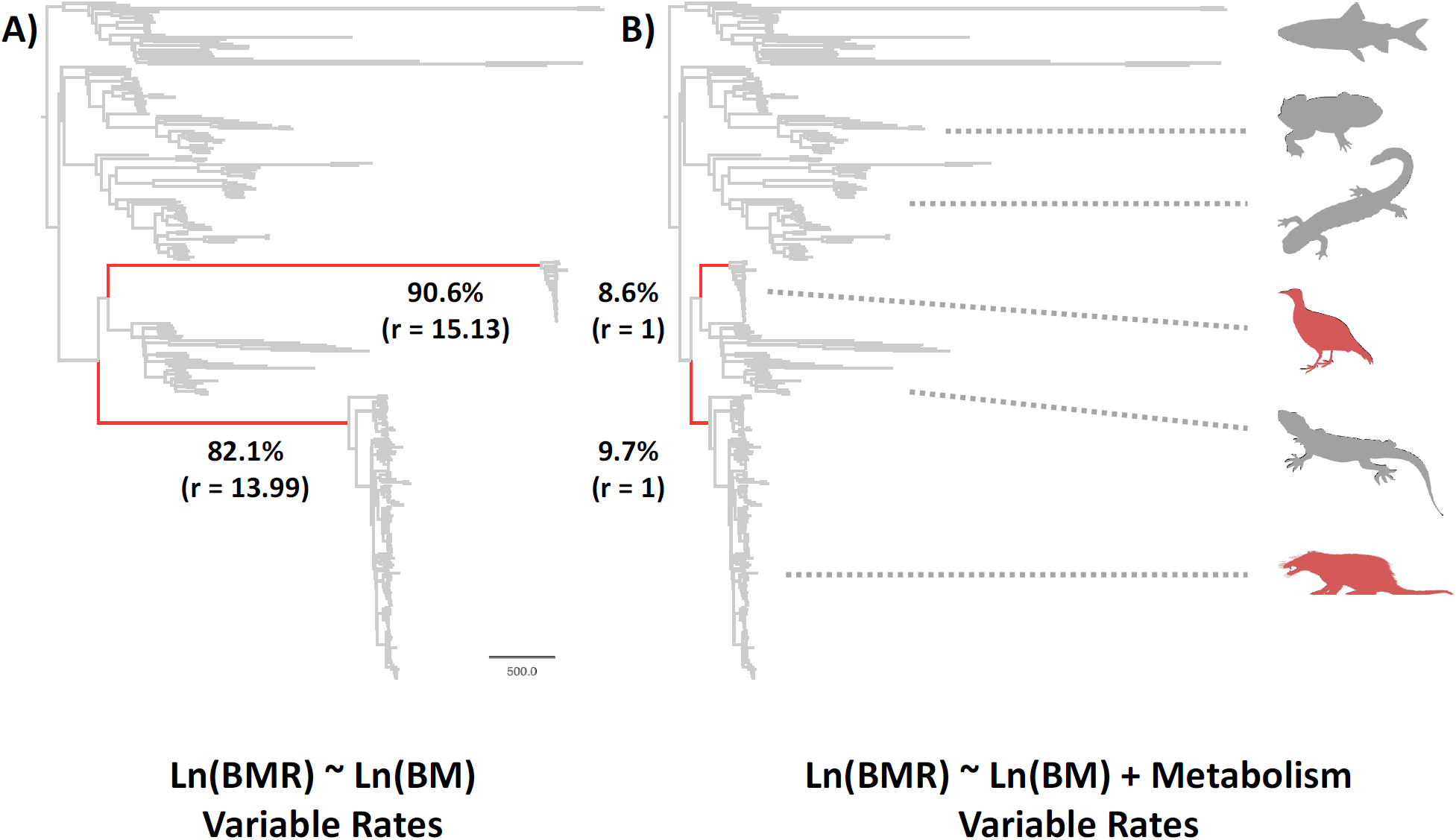
Rate-scaled trees from the variable rates regression analyses without (A) and with (B) the presence of endothermy. The most-supported model (B) allows for a difference in BMR between endotherms and ectotherms, accounting for body mass (BIC = 814.64). The null model (A) ranks third (ΔBIC = 17.03). The two red-colored branches lead to the common ancestors of birds and mammals. The bold percentages represent the percent of the posterior distribution of which the red-colored branches were positively scaled (r > 1). These percentages are considerably reduced when we allow for a difference in BMR between endotherms and ectotherms (90.6 to 8.6% for birds and 82.1 to 9.7% for mammals); median rate scalars are also greatly reduced (15.13 and 13.99 to 1 for birds and mammals, respectively). Silhouettes were sourced from phylopic.org: Aves by George Edward Lodge (modified by T. Michael Keesey); Cryptobranchoidea by zoosnow; Lepidosauria by Ghedo and T. Michael Keesey; Mammalia by T. Michael Keesey; Salientia by Nobu Tamura; *Salvelinus namaycush* by Sherman F. Denton via rawpixel.com (illustration) and Timothy J. Bartley (silhouette).

We found no support for a punctuated model of BMR evolution. There is little to no evidence for net speciation having a substantial effect on the rate of BMR evolution (pMCMC = 0.37). BMR did not evolve more rapidly along extant vertebrate lineages that speciated more frequently. We additionally found no evidence for an effect of net speciation on ln BMR, conditioning on ln body mass and the presence/absence of endothermy (pMCMC = 0.41). This suggests that the frequency of speciation along vertebrate lineages cannot explain the variation we observe in BMR. It also implies that speciation had no consistent directional effects on BMR evolution—more frequent speciation did not result in lower or higher metabolic rates. Net speciation did not have an effect on the rate of BMR evolution or the variation in BMR that manifested in living vertebrates. The results from our full dataset were corroborated by the analyses using our three randomly down-sampled datasets. The only difference was that, for two of our down-sampled datasets, a variable rates model of BMR evolution was not supported over one that assumes a single uniform rate (BF > 5.0). Since this results in all taxa having equal path lengths regardless of node count, there is little evidence that net speciation had an effect on the rate of BMR evolution for these down-sampled datasets (random samples 1 and 2). For the third down-sampled dataset, there was evidence for variable rates of BMR evolution (BF = 5.78) but little evidence for an effect of net speciation on the rate of BMR evolution (pMCMC > 0.05). We also found no evidence for the effect of net speciation on ln BMR variation, conditioning on ln body mass and the presence/absence of endothermy, for the first two randomly down-sampled datasets (pMCMC = 0.65 and 0.76). The third down-sampled dataset yielded statistical significance, but the effect of net speciation on ln BMR was negligible (pMCMC = 0.046, slope = 0.043). The taxon lists for each of the three down-sampled datasets can be found in Table S7.

## Discussion

Despite theoretical expectations, there is ambiguous support for a link between metabolic rate and genome size in the literature. A previous summary of research suggests that there may be a difference in the effect of genome size between endothermic and non-endothermic vertebrates with consistent evidence for an effect in mammals but not in actinopterygians or lissamphibians [3]. There is ambiguous support for an effect in birds, which may be due to differences in sample size and variation among methods. Our study, which leverages phylogenetic comparative methods, aims to test for the relationship between BMR and genome size in extant vertebrates while accounting for variable rates of evolution. We successfully replicated Uyeda et al.’s results [9] in which the effect of genome size was not included in our final model. This result is inconsistent with previous studies on amniotes that found evidence for an effect, particularly in birds [5–7], though others did not find evidence for a relationship [4,8]. Given previous suggestions in the literature, we further included an interaction between genome size and the presence/absence of endothermy to test for a difference in the effect of genome size on BMR; this parameter was not included in our final model either. These results indicate, despite previous suggestions, that differences in the effect of genome size among metabolic strategies does not sufficiently improve models in explaining metabolic rate variation. This is consistent with our variable rates regression results in which we find no evidence that genome size explains the rate variation observed in BMR after accounting for body mass. We do find some evidence that the presence of endothermy explains evolutionary rate shifts in BMR along branches leading to birds and mammals, but differences in metabolic strategies do not explain rate variation globally. It is likely that other variables, such as life history traits linked to body mass, overshadow any association between genome size and BMR—as suggested by Vinogradov [4]. Regardless, genome size is thought to be indirectly linked to BMR by way of its influence on nucleus and cell size [6]. Moreover, this relationship may vary by clade owing to underlying differences in biology. Uyeda et al. [9] find support for these nuances. They demonstrate that major shifts in metabolic rate cannot be explained by genome size alone, despite some evidence for the latter explaining variation in metabolic rate. However, testing this would require denser sampling within each vertebrate clade. Larger sample sizes and more complex phylogenetic models may yield evidence that genome size influences metabolic rate variation within certain clades, as in Gregory’s [6] larger sample of birds. Differences in the tempo and mode of genome size and metabolic evolution may also obscure functional relationships between the two—as previously suggested by Waltari and Edwards [7] and Uyeda et al. [9]. We find no evidence for a punctuated mode of BMR evolution; specifically, there is little to no evidence for an effect of speciation on either the rate of BMR evolution or the observed variance in BMR.

Further studies on the relationship between metabolism and genome size will benefit from denser sampling across clades, sophisticated phylogenetic comparative methods, and more experimental studies, which together will help clarify how genome organization may relate to physiology in other groups of vertebrates. Moreover, our review briefly highlights the importance of analyzing other types of bone histological data, beyond osteocyte lacunae size (e.g., vascularization, tissue type, etc.), in paleophysiological studies [28]. These data can provide further insight into the associations between bone growth and metabolism, and help uncover the deep evolutionary history of animal physiology.

## Acknowledgments

We thank Jorge Cubo and Adam Huttlenlocker for inviting us to contribute to this special issue and Kevin Surya for discussions about this paper. We also thank Alexander Suh for comments on an early version of this paper as well as Michael D’Emic and two anonymous reviewers for improving the manuscript.

