## Supplementary Material for "The Relationship Between Genome Size and Metabolic Rate in Extant Vertebrates"

**<u>Supplementary Materials</u>**

**Table of Contents** **Page**

Supplementary tables
Table 1: Variance inflation factors for final model 2
Table 2: Model log marginal likelihoods and BIC values 2
Table 3: BF comparisons between VR and uniform-rate models 3
Table 4: BIC comparisons among VR models 3
Table 5: Parameter estimates for final model 3
Table 6: Taxa with highly positive rate shifts in final model 4
Table 7: Taxa lists for three randomly down-sampled datasets 5

Supplementary figures
Figure S1: Residual vs fitted plot for final model 7
Figure S2: Q-Q plot for final model 8
Figure S3: Scale-location plot for final model 9
Figure S4: Residuals vs leverage plot for final model 10
Figure S5: Likelihood trace plot for final uniform-rate model 11
Figure S6: Likelihood trace plot for final VR model 12

References cited 12

**Table 1**: Variance inflation factors for chosen model variables without variable rates (Model Nodes).

vif(Model.3): lnBMR ~ lnMass + endothermy

| lnMass | endo |
|---|---|
| 1.244792 | 1.244792 |

The variance inflation factor (VIF) is a measure of multicollinearity among variables in a multiple regression model. It measures the ratio of the full model's variance over the variance of a model with only a single explanatory variable (headers indicate chosen variable). VIF values over 5 indicate evidence for multicollinearity. The two variables displaying the highest VIF values indicate which two variables are collinear. We find no evidence for multicollinearity in a multiple regression model that includes our chosen variables, assuming uniform rates. We used the vif function in the R package, *car*, to estimate the VIF values [1]. We used a linear model in R to assess model assumptions of equal-variance and residual normality (see Supp. Figs. 1-4).

**Table 2**: Log-marginal likelihoods (Lh) and Bayesian Information Criterion (BIC) values for the models analyzed. The number of parameters = # $\beta$ coefficients + $\alpha$ + median # branch- and clade-specific rate scalars. The most-supported model is indicated in bold. VR = variable rates.

| Model ID | Model Description | Log Marginal Lh | # Parameters | BIC |
|---|---|---|---|---|
| Null | LnBMR ~ LnBM | -327.032476 | 2 | 665.5827555 |
| Null-VR | - | -271.862481 | 50 | 831.6700507 |
| A1 | LnBMR ~ LnBM + LnGS | -331.847033 | 3 | 680.9707713 |
| A1-VR | - | -275.174193 | 52 | 849.8112782 |
| A2 | LnBMR ~ LnBM + LnGS + endothermy | -329.869751 | 4 | 682.7751091 |
| A2-VR | - | -267.249539 | 51 | 828.2030685 |
| A3 | LnBMR ~ LnBM + endothermy | -324.575411 | 3 | 666.4275273 |
| **A3-VR** | **-** | **-263.349659** | **50** | **814.6444067** |
| Full | LnBMR ~ LnBM + LnGS + endothermy + LnGS*endothermy | -335.209148 | 5 | 699.2128049 |
| Full-VR | - | -273.937544 | 52 | 847.3379802 |

**Table 3**: Bayes factors (BF) comparing variable rates models (VR) with uniform-rate models. We found evidence for a VR model of evolution with all regression models (BF > 2).

| Model Comparison | Bayes Factor |
|---|---|
| **Null-VR vs Null** | 110.33999 |
| **A1-VR vs A1** | 113.34568 |
| **A2-VR vs A2** | 125.240424 |
| **A3-VR vs A3** | 122.451504 |
| **Full-VR vs Full** | 122.543208 |

**Table 4**: BIC model comparisons for VR regression models. The model with the lowest BIC value is the top model. Any model comparisons with a difference in BIC value (ΔBIC) over 2 is evidence for a significant difference in favor of the top model. Model A3-VR was our top model.

| Model Comparison | BIC | ΔBIC |
|---|---|---|
| A3-VR | 814.6444067 | - |
| A2-VR | 828.2030685 | 13.55866177 |
| Null-VR | 831.6700507 | 17.025644 |
| Full-VR | 847.3379802 | 32.69357355 |
| A1-VR | 849.8112782 | 35.16687155 |

**Table 5**: Estimated parameters for Model A3-VR.

| Model Parameter | Value |
|---|---|
| Avg. $\alpha$ | -2.128338484 |
| Avg. $\beta_1$ (LnBM) | 0.733004854 |
| Avg. $\beta_2$ (Endothermy) | 4.265239484 |
| Avg. Var | 0.002485956 |
| Avg. $R^2$ | 0.87471748 |
| Avg. s.e. $\alpha$ | N/A |
| Avg. s.e. $\beta_1$ | 0.015960608 |
| Avg. s.e. $\beta_2$ | 0.633467364 |
| Median RJ Local Branch | 30 |
| Median RJ Local Node | 17 |

**Table 6:** Taxa with highly positive rate shifts in the final model, including the node identifier, the branch length from the original time tree, the percent time it was scaled in the posterior distribution of models, the median scalar of the posterior distribution of scaled branches, and the terminal taxo located on the tip of each scaled branch.

| Node ID | Original BL | Pct time scaled | Median Scalar | Taxa List |
|---|---|---|---|---|
| **59** | 2.41583 | 99.973333 | 20.669061 | *Pteropus_vampyrus* |
| **351** | 22.097213 | 99.893333 | 10.800664 | *Lacerta_viridis* |
| **368** | 54.722746 | 98.6 | 9.804216 | *Sceloporus_occidentalis* |
| **413** | 10.978897 | 96.066667 | 5.50693 | *Ambystoma_mexicanum, Ambystoma_tigrinum* |
| **414** | 5.817084 | 99.773333 | 8.068862 | *Ambystoma_tigrinum* |
| **415** | 5.817084 | 96.093333 | 5.82247 | *Ambystoma_mexicanum* |
| **417** | 2.917064 | 95.066667 | 4.551631 | *Ambystoma_macrodactylum, Ambystoma_talpoideum* |
| **418** | 13.862921 | 95.306667 | 4.564875 | *Ambystoma_talpoideum* |
| **420** | 1.893286 | 95.373333 | 4.623398 | *Ambystoma_jeffersonianum, Ambystoma_maculatum* |
| **421** | 14.8867 | 95.84 | 4.671601 | *Ambystoma_maculatum* |
| **422** | 14.8867 | 95.213333 | 4.590435 | *Ambystoma_jeffersonianum* |
| **425** | 7.730314 | 99.426667 | 28.380807 | *Salamandra_infraimmaculata* |
| **458** | 0.96325 | 98.053333 | 8.261879 | *Desmognathus_fuscus, Desmognathus_ochrophaeus* |
| **459** | 12.792203 | 96.493333 | 6.607416 | *Desmognathus_fuscus* |
| **460** | 12.792203 | 96.92 | 7.042009 | *Desmognathus_ochrophaeus* |
| **477** | 20.104453 | 99.92 | 14.406443 | *Eurycea_nana, Eurycea_neotenes* |
| **478** | 1.14061 | 99.293333 | 12.893374 | *Eurycea_nana* |
| **479** | 1.14061 | 99.666667 | 14.01603 | *Eurycea_neotenes* |
| **549** | 16.108825 | 95.96 | 6.38115 | *Pseudacris_nigrita, Pseudacris_triseriata* |
| **550** | 4.752883 | 99.746667 | 8.13352 | *Pseudacris_nigrita* |
| **551** | 4.752883 | 96.16 | 6.647734 | *Pseudacris_triseriata* |
| **554** | 16.586995 | 96.693333 | 5.765494 | *Hyla_squirella* |
| **567** | 29.00934 | 96.293333 | 7.723469 | *Bufo_alvarius, Bufo_americanus, Bufo_terrestris, Bufo_viridis* |
| **568** | 3.066473 | 98.293333 | 10.094302 | *Bufo_alvarius, Bufo_americanus, Bufo_terrestris* |
| **569** | 23.238186 | 98.2 | 9.829487 | *Bufo_alvarius* |
| **570** | 20.450294 | 98.786667 | 11.979241 | *Bufo_americanus, Bufo_terrestris* |
| **571** | 2.78789 | 99.893333 | 19.754249 | *Bufo_terrestris* |
| **572** | 2.78789 | 98.906667 | 14.454481 | *Bufo_americanus* |

| | | | | |
|---|---|---|---|---|
| **573** | 26.30466 | 96.76 | 7.999324 | *Bufo_viridis* |
| **589** | 20.96477 | 95.16 | 6.511558 | *Lepomis_gibbosus* |
| **604** | 96.479581 | 95.626667 | 14.387232 | *Hippoglossoides_platessoides, Pseudopleuronectes_americanus* |
| **605** | 12.05258 | 99.973333 | 36.242105 | *Hippoglossoides_platessoides* |
| **606** | 12.052582 | 95.72 | 18.004253 | *Pseudopleuronectes_americanus* |
| **608** | 117.909691 | 98.706667 | 8.525924 | *Typhlogobius_californiensis* |

**Table 7:** Taxa list for three randomly down-sampled datasets.

| Dataset | Taxa List (in the following order: Anura, Lepidosauria, Urodela, Aves, and Mammalia) |
|---|---|
| **Random 1** | Discoglossus_pictus, Leptodactylus_fuscus, Pseudacris_crucifer, Rana_esculenta, Scaphiopus_holbrookii, Rana_arvalis, Pyxicephalus_adspersus, Bombina_orientalis, Hyperolius_marmoratus, Rana_temporaria, Hyla_gratiosa, Rana_sylvatica, Hyla_chrysoscelis, Pseudacris_regilla, Mannophryne_trinitatis, Spea_bombifrons, Rana_palustris, Rana_ridibunda, Crinia_signifera, Iguana_iguana, Sceloporus_occidentalis, Lacerta_agilis, Lampropeltis_getula, Psammodromus_algirus, Epicrates_cenchria, Liophis_miliaris, Waglerophis_merremi, Vipera_berus, Sphenodon_punctatus, Trogonophis_wiegmanni, Phrynosoma_cornutum, Boa_constrictor, Podarcis_muralis, Varanus_bengalensis, Chalcides_ocellatus, Elgaria_multicarinata, Coluber_constrictor, Anolis_carolinensis, Gekko_gecko, Natrix_maura, Hemidactylus_frenatus, Thamnophis_sirtalis, Anguis_fragilis, Ptyodactylus_hasselquistii, Natrix_natrix, Python_curtus, Eunectes_murinus, Plethodon_cinereus, Pseudoeurycea_rex, Coereba_flaveola, Melospiza_georgiana, Colius_colius, Larus_argentatus, Sayornis_phoebe, Geothlypis_trichas, Falco_sparverius, Melospiza_melodia, Phalacrocorax_auritus, Dendroica_palmarum, Dendroica_dominica, Parus_atricapillus, Cardinalis_cardinalis, Seiurus_aurocapilla, Coturnix_coturnix, Zonotrichia_albicollis, Dendroica_coronata, Bonasa_umbellus, Myiarchus_crinitus, Carduelis_tristis, Zonotrichia_leucophrys, Ammodramus_savannarum, Seiurus_noveboracensis, Agelaius_phoeniceus, Spizella_passerina, Pica_pica, Icterus_galbula, Passerculus_sandwichensis, Contopus_virens, Dipodomys_agilis, Thomomys_bottae, Uroderma_bilobatum, Pteropus_poliocephalus, Eumops_perotis, Cavia_porcellus, Procavia_capensis, Setifer_setosus, Pteropus_rodricensis, Heteromys_anomalus, Eulemur_fulvus, Heliophobius_argenteocinereus, Ctenomys_opimus, Microtus_californicus, Lagostomus_maximus, Zaedyus_pichiy, Pteronotus_parnellii |
| **Random 2** | Acris_crepitans, Bufo_viridis, Rana_arvalis, Leptodactylus_fuscus, Rana_esculenta, Odontophrynus_americanus, Pyxicephalus_adspersus, Osteopilus_septentrionalis, Xenopus_muelleri, Pseudacris_crucifer, Rana_sylvatica, Rana_erythraea, Mannophryne_trinitatis, Scaphiopus_couchii, Rana_palustris, Spea_bombifrons, Scaphiopus_holbrookii, Xenopus_laevis, Hyla_squirella, Eunectes_murinus, Tarentola_mauritanica, Lacerta_viridis, Vipera_berus, Psammodromus_algirus, Lacerta_agilis, Anguis_fragilis, Epicrates_cenchria, Python_curtus, Coluber_constrictor, Chalcides_ocellatus, Gekko_gecko, Coleonyx_variegatus, Acanthodactylus_pardalis, Waglerophis_merremi, |

| | |
|---|---|
| | Iguana_iguana, Ptyodactylus_hasselquistii, Natrix_maura, Lampropeltis_getula, Elgaria_multicarinata, Varanus_bengalensis, Tiliqua_scincoides, Natrix_natrix, Anolis_carolinensis, Crotaphytus_collaris, Boa_constrictor, Podarcis_muralis, Hemidactylus_frenatus, Plethodon_cinereus, Taricha_torosa, Coereba_flaveola, Melospiza_georgiana, Colius_colius, Larus_argentatus, Sayornis_phoebe, Geothlypis_trichas, Falco_sparverius, Melospiza_melodia, Phalacrocorax_auritus, Dendroica_palmarum, Dendroica_dominica, Parus_atricapillus, Cardinalis_cardinalis, Seiurus_aurocapilla, Coturnix_coturnix, Zonotrichia_albicollis, Dendroica_coronata, Bonasa_umbellus, Myiarchus_crinitus, Carduelis_tristis, Zonotrichia_leucophrys, Ammodramus_savannarum, Seiurus_noveboracensis, Agelaius_phoeniceus, Spizella_passerina, Pica_pica, Icterus_galbula, Passerculus_sandwichensis, Contopus_virens, Heteromys_anomalus, Thomomys_talpoides, Dobsonia_moluccensis, Erinaceus_europaeus, Lasiorhinus_latifrons, Apodemus_sylvaticus, Macroglossus_minimus, Rousettus_amplexicaudatus, Chaetophractus_villosus, Mormoops_megalophylla, Noctilio_albiventris, Canis_latrans, Perameles_gunnii, Potorous_tridactylus, Trichosurus_vulpecula, Sturnira_lilium, Peromyscus_crinitus |
| **Random 3** | Spea_hammondii, Leptodactylus_fuscus, Gastrophryne_carolinensis, Rana_temporaria, Odontophrynus_americanus, Rana_esculenta, Pseudacris_nigrita, Crinia_signifera, Pseudacris_triseriata, Pyxicephalus_adspersus, Xenopus_laevis, Pseudacris_regilla, Hyla_cinerea, Scaphiopus_holbrookii, Mannophryne_trinitatis, Bufo_terrestris, Acris_crepitans, Bufo_americanus, Kassina_senegalensis, Phrynosoma_cornutum, Python_curtus, Waglerophis_merremi, Natrix_natrix, Thamnophis_sirtalis, Lacerta_agilis, Psammodromus_algirus, Trogonophis_wiegmanni, Lacerta_viridis, Sphenodon_punctatus, Varanus_bengalensis, Gekko_gecko, Anolis_carolinensis, Eunectes_murinus, Tiliqua_scincoides, Epicrates_cenchria, Boa_constrictor, Acanthodactylus_pardalis, Vipera_berus, Hemidactylus_frenatus, Anguis_fragilis, Lampropeltis_getula, Iguana_iguana, Elgaria_multicarinata, Crotaphytus_collaris, Coleonyx_variegatus, Chalcides_ocellatus, Coluber_constrictor, Aneides_hardii, Plethodon_glutinosus, Coereba_flaveola, Melospiza_georgiana, Colius_colius, Larus_argentatus, Sayornis_phoebe, Geothlypis_trichas, Falco_sparverius, Melospiza_melodia, Phalacrocorax_auritus, Dendroica_palmarum, Dendroica_dominica, Parus_atricapillus, Cardinalis_cardinalis, Seiurus_aurocapilla, Coturnix_coturnix, Zonotrichia_albicollis, Dendroica_coronata, Bonasa_umbellus, Myiarchus_crinitus, Carduelis_tristis, Zonotrichia_leucophrys, Ammodramus_savannarum, Seiurus_noveboracensis, Agelaius_phoeniceus, Spizella_passerina, Pica_pica, Icterus_galbula, Passerculus_sandwichensis, Contopus_virens, Elephantulus_edwardii, Procyon_lotor, Tympanoctomys_barrerae, Perameles_nasuta, Geomys_bursarius, Myrmecophaga_tridactyla, Glossophaga_soricina, Dipodomys_agilis, Pteronotus_parnellii, Perodicticus_potto, Microtus_californicus, Chalinolobus_gouldii, Dipodomys_panamintinus, Didelphis_marsupialis, Phascolarctos_cinereus, Sigmodon_hispidus, Saimiri_sciureus |

**Figure 1**: Residuals of lnBMR vs fitted plot for final model (Model A3). The distribution of the residuals against the fitted values is used to assess the assumption of equal variance in the residual error. There is a violation in equal-variance due to increasing residual error in moderate fitted values. However, the residual error does not match that of the final model assessed because we did not use phylogenetic comparative methods or account variable rates of evolution in making this plot.

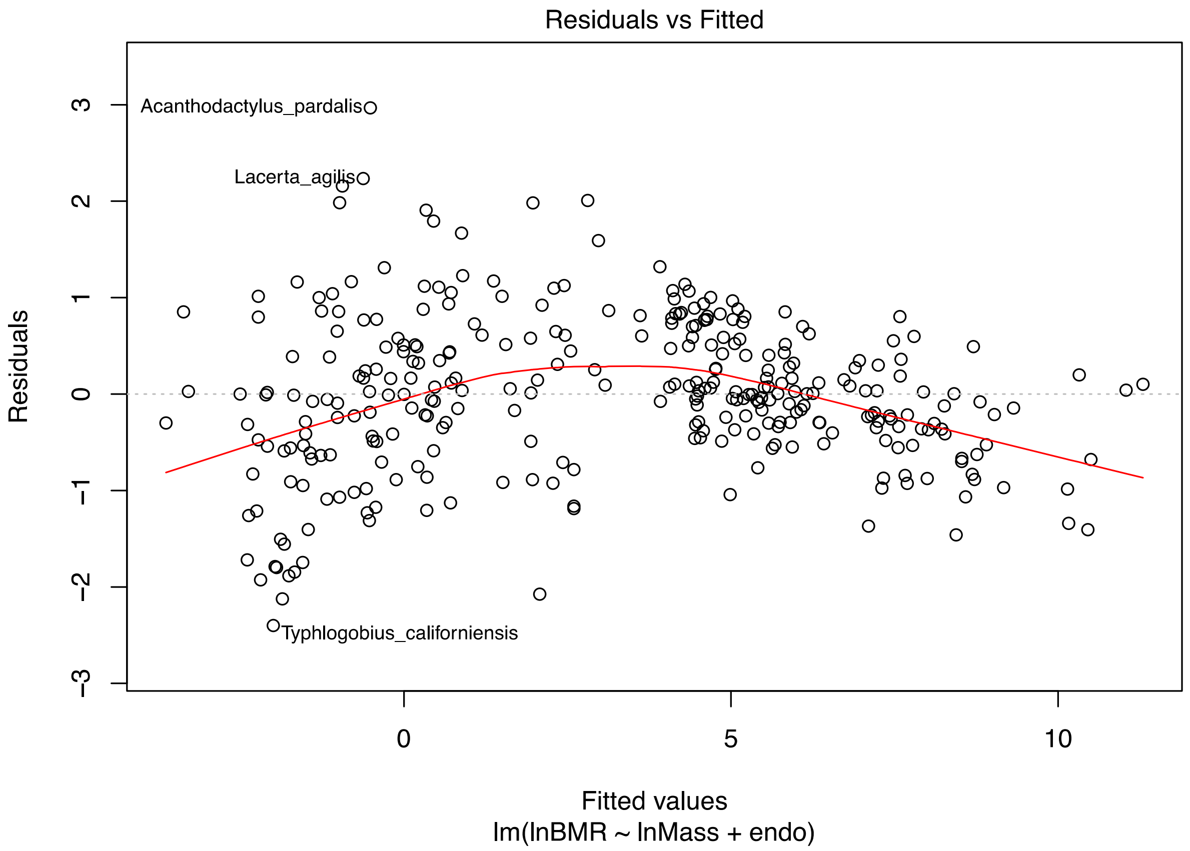

**Figure 2**: Q-Q plot of the residuals from the final model (Model A3), used to evaluate the assumption that residual errors are normally distributed. This There is slight deviation from residual normality due to a thick-tail effect; however, as previously mentioned, these residuals are not entirely accurate because we did not use phylogenetic comparative methods or account for variable rates of evolution when making this plot.

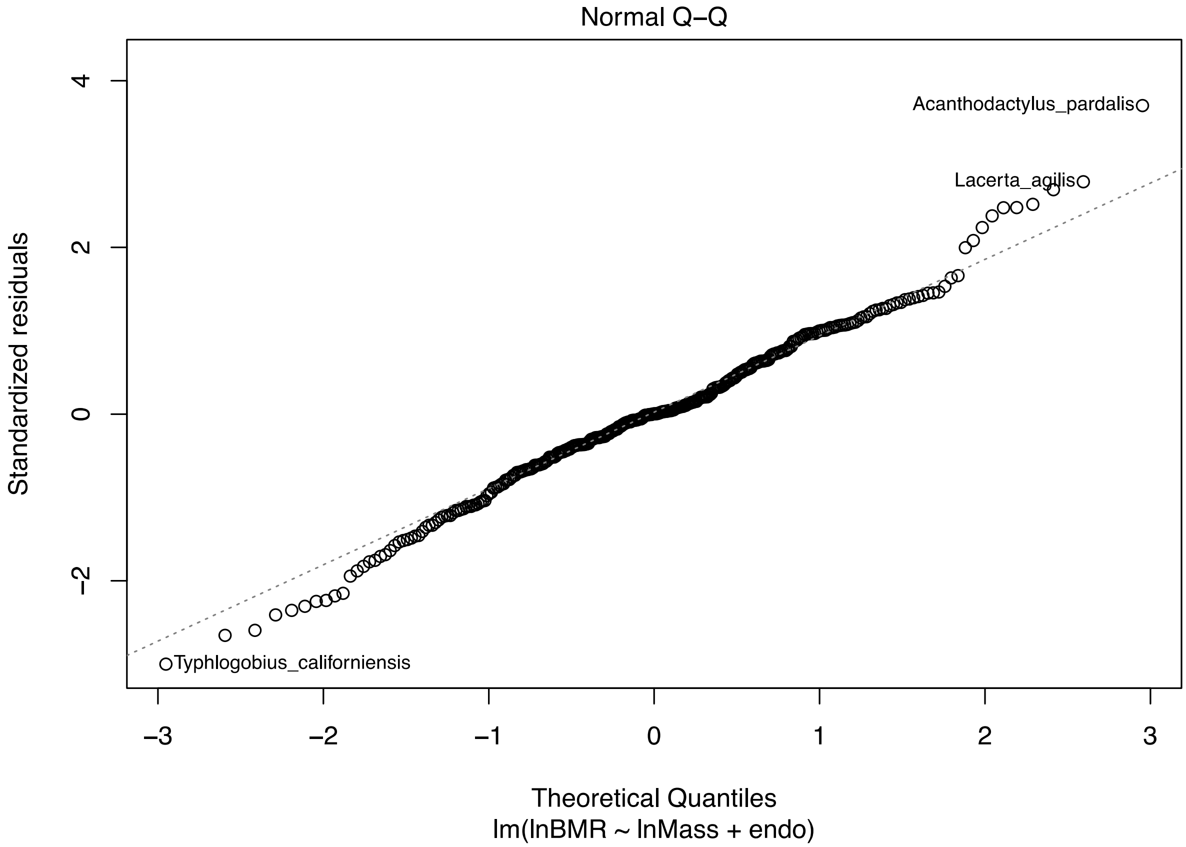

**Figure 3**: Scale-location plot for the final model, used to evaluate the assumption of equal variance in the standardized residual errors. There is a slight violation in the assumption of equal variance with a decrease in residual variance as fitted values increase.

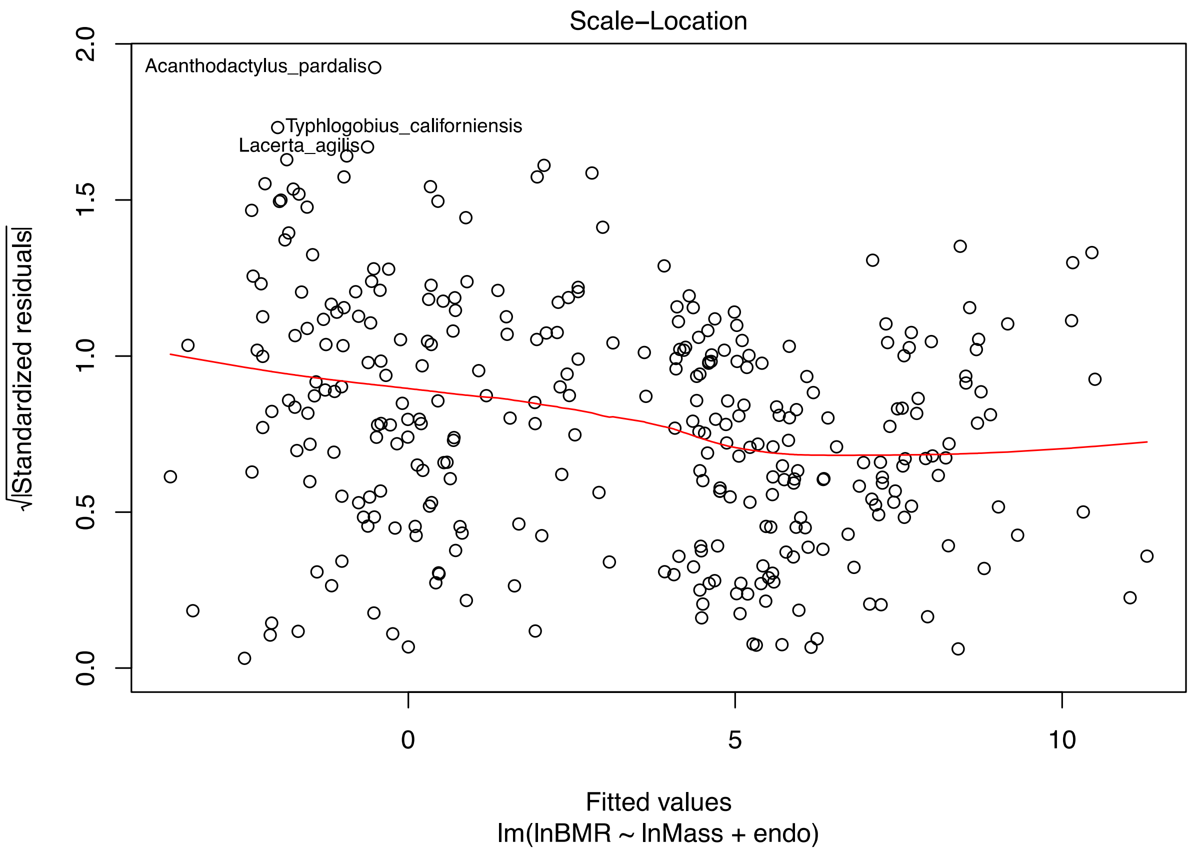

**Figure 4**: Residual vs leverage plot for the final model, used for evaluating the leverage of each data point on regression model parameter estimates. Data with more extreme residual errors can have an influence (i.e., leverage) on the model estimates, such as the slope of the line. Although a few data points have higher leverage than others, these points do not have a large Cook's distance, indicating that none of the data points exhibit high leverage on the parameter estimates.

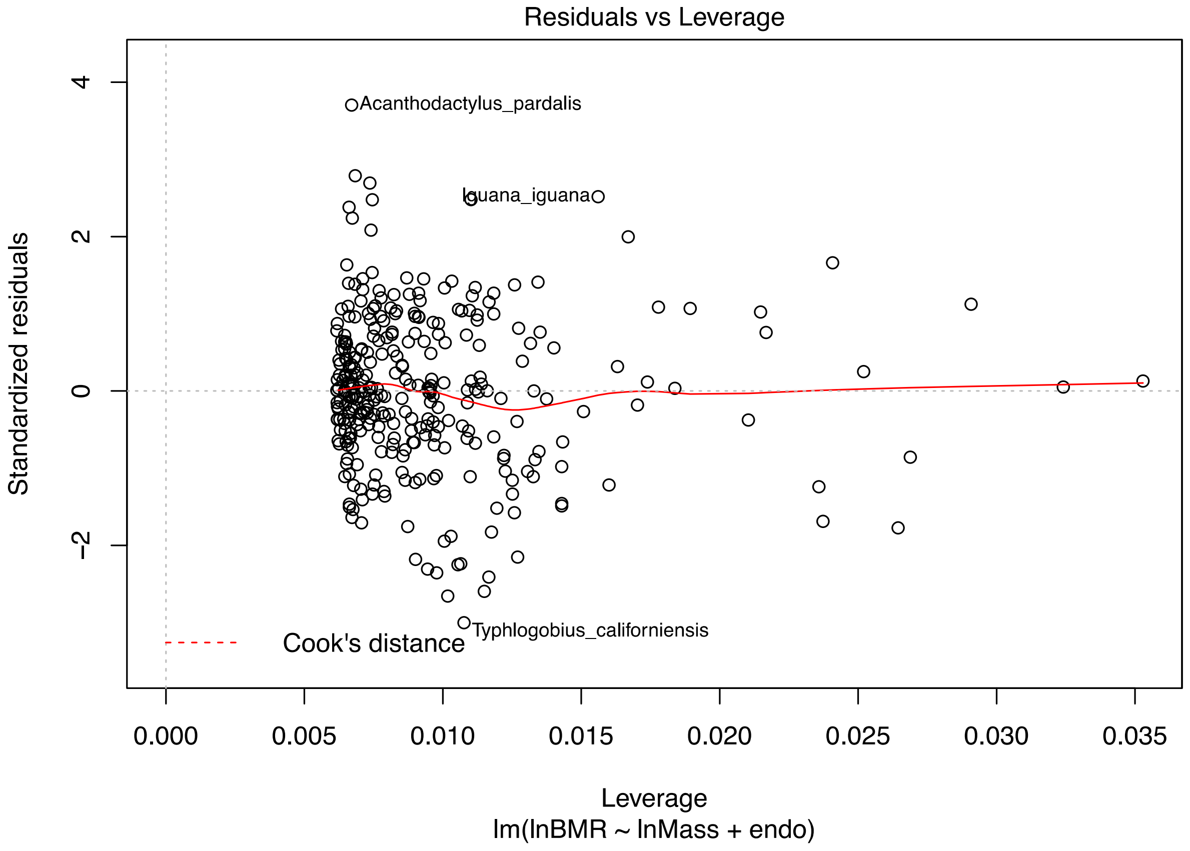

**Figure 5**: Likelihood trace plot of the final uniform-rate model (Model A3). Trace plot was made using the program, Tracer 1.7 [2]. The burn-in is represented by the transparent region of the plot. Effective sample size = 6500.

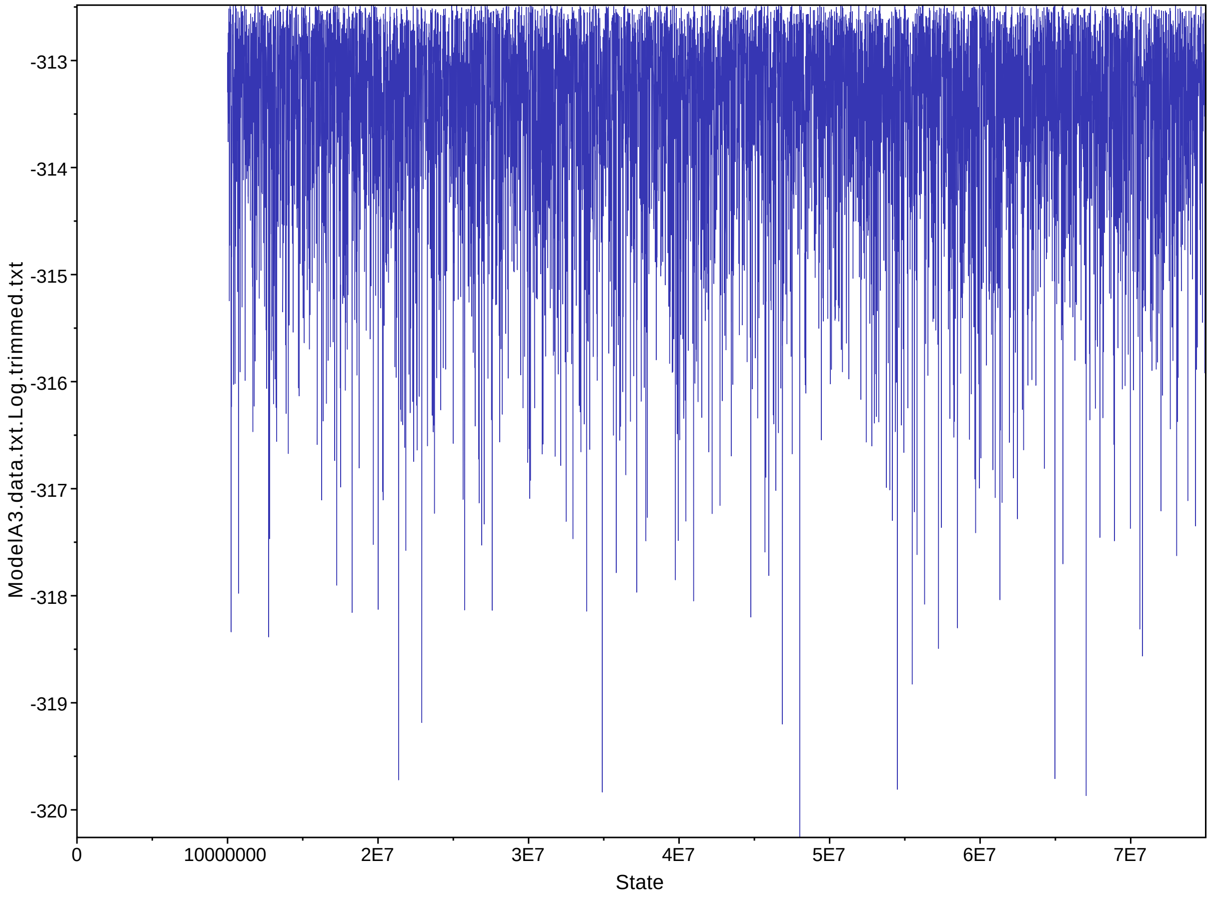

**Figure 6**: Likelihood trace plot of the final variable rates model (Model A3-VR). Trace plot was made using the program, Tracer 1.7 [2]. The burn-in is represented by the transparent region of the plot. Effective sample size = 4356.

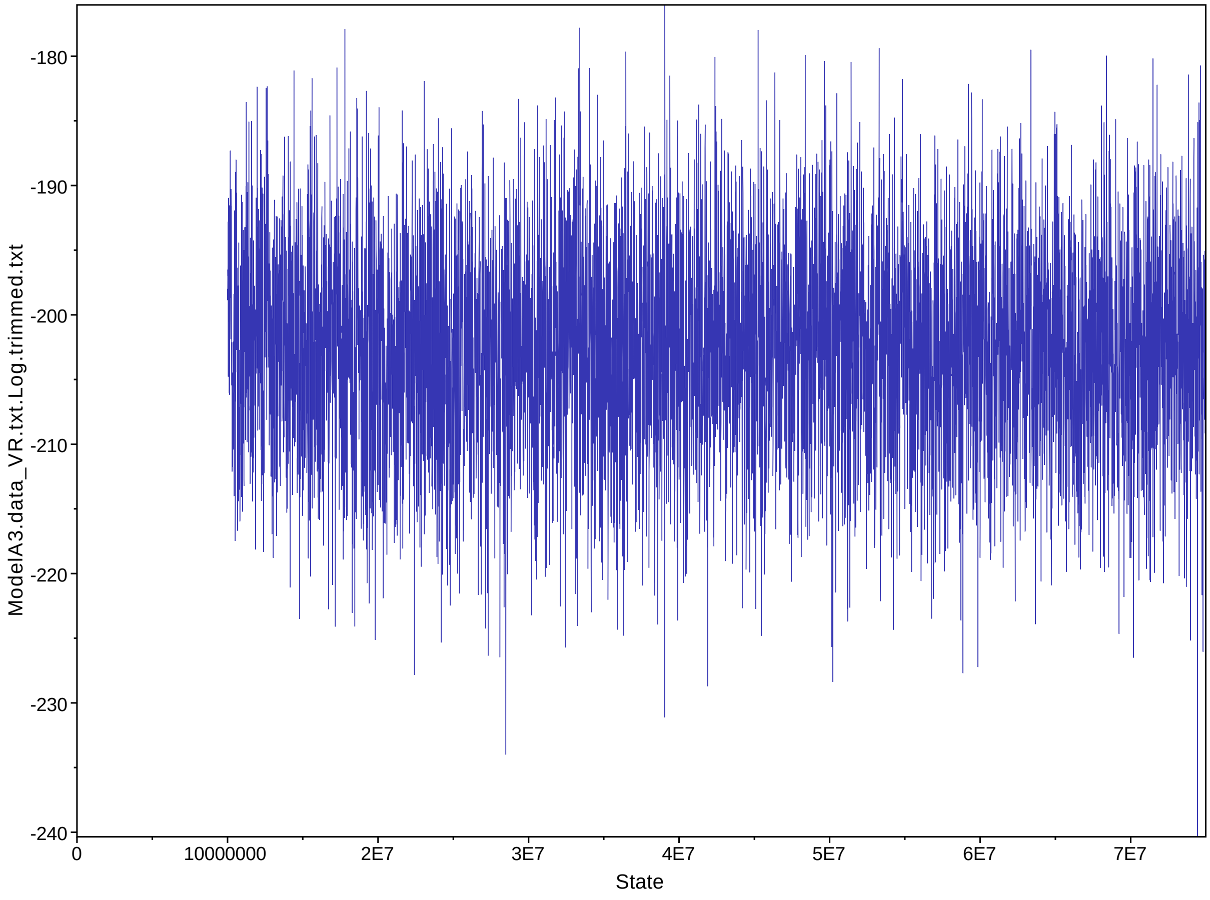
